# Inheritance of somatic mutations can affect fitness in monkeyflowers

**DOI:** 10.1101/2024.05.20.595007

**Authors:** Matthew A. Streisfeld, Jessie C. Crown, Jack J. McLean, Aidan W. Short, Mitchell B. Cruzan

## Abstract

Plants possess the unique ability to transmit mutations to progeny that arise both through meiotic and mitotic (somatic) cell divisions. This is because the same meristem cells responsible for vegetative growth also generate gametes for sexual reproduction. Despite the potential for somatic mutations to be an additional source of genetic variation for adaptation, their role in plant evolution remains largely unexplored. We performed multiple experiments in the bush monkeyflower (*Mimulus aurantiacus*) to determine the fitness effects of somatic mutations inherited across generations. We tracked somatic mutations transmitted to progeny by generating self-pollinations within a flower (autogamy) or between stems of the same plant (geitonogamy). Autogamy and geitonogamy lead to different segregation patterns of somatic mutations among stems, making it possible to compare average fitness due to somatic variants. We found increased fecundity following autogamy, as well as significant impacts on drought tolerance, survival, and biomass. The variance in fitness was also greater following autogamy, consistent with the effects of somatic mutations impacting fitness. Effect sizes were small, but predictable, given that *M. aurantiacus* is a long-lived, drought-adapted shrub. These results reveal the importance of inherited somatic mutations as a source of genetic variation that can be relevant for plant adaptation.

## Introduction

Mutations are the ultimate source of genetic variation. While this is a well-known saying in genetics, only mutations that are transmitted to subsequent generations will be relevant for evolution. Mutations are generated through both mitosis and meiosis, but among most animals, only mutations that arise in the germline can be transmitted to progeny. This is because the germline is determined early in development and is separated from the somatic cell lineages that form the rest of the organism’s body. Therefore, the set of somatic mutations that form during mitotic division outside of the germline are typically not heritable.

By contrast, plants undergo indeterminate growth, where shoot and root systems continually elongate and develop throughout a significant portion of their life-cycle (Antolin & Strobeck. 1985, D’Amato 1996). Growth of the shoot system in plants occurs at shoot apical meristems (SAMs), which contain a population of undifferentiated cells known as the central zone. In vascular plants, these cells differentiate into leaf and stem tissue necessary for growth and development, and they eventually produce the gametes required for sexual reproduction. This reservoir of pluripotent cells is continually replenished through mitotic division (Kwiatkowska 2008), but as the shoot elongates, somatic mutations may occur due to DNA replication errors. These somatic mutations can accumulate as the stem elongates, resulting in distal areas of the shoot system possessing more somatic variants than their basal counterparts (Schultz & Scofield 2009). In angiosperms, the gametes are not produced until later in development when the SAM is first converted to a floral meristem and then to a flower, indicating that somatic mutations may be transmitted to offspring. This leads to the possibility that somatic mutations are an important source of genetic variation that can impact evolutionary processes.

Despite the potential for the inheritance of somatic variants that accumulated during vegetative growth, the role and relevance of somatic mutations within plants remains unsettled. Since plants possess the ability to pass on both meiotic and somatic mutations to progeny, one might expect the mutation rate per generation among plants would be noticeably higher than animals. However, mutation rates per generation appear to be similar between plants and animals (Gaut et al. 2011). Multiple explanations have been offered to explain this discrepancy.

For example, germline segregation in plants may occur earlier in development than previously appreciated, with primordial germ cells physically separated from future somatic cells within the meristem (Lanfear et al. 2018). This explanation asserts that somatic mutations arising during vegetative growth are only rarely inherited by progeny, since future germ cells would only be found in isolated cell lineages (Cruzan 2018). These isolated populations of germ cells could potentially have a slower rate of division than their somatic counterparts, and as a result, they would have a significantly lower mutation rate over time (Lanfear et al. 2018). Due to its slow cell division rate relative to the peripheral zone and rib meristem, the central zone of the angiosperm SAM is a candidate location for this proposed population of segregated germ cells (Cutter 1965). However, additional observations in angiosperm models have revealed that mitotic activity spikes both within the central zone and rib meristem during the transition from vegetative to reproductive tissue, suggesting that multiple zones contribute to the formation of the gametes (Kwiatkowska 2008). More recently, computational models based on quantitative cell lineage data from *Arabidopsis thaliana* and tomato (*Solanum lycopersicum*) were used to replicate patterns of cell division in SAMs and axillary meristems (Burian et al. 2016). These models suggested that cells were not constantly replaced within the central zone of the SAM, and instead persisted throughout vegetative growth. Burian et al. (2016) claimed these findings indicated that plants possess mechanisms to prevent the fixation and eventual accumulation of deleterious genetic load. They further asserted that plants possess germlines analogous to those found within animals.

An alternate explanation posits that cell lineages containing deleterious somatic mutations are removed from the population of meristem cells due to natural selection (Cruzan 2018). This has been referred to as cell lineage selection (CLS; Fagerstrom et al. 1998; Otto and Hastings 1998; Monro and Poore 2009). Since the size of the central zone is fixed and is constantly replenished through mitotic division, cell lineages that express deleterious mutations may replicate more slowly and therefore will be replaced by cell lineages with accelerated division (Pineda-Krch & Lehtila 2002). Models of stochastic growth have indicated that relatively minor differences in cell replication rates during development can result in significant differences in the proportion of mutant cells found within adults (Otto & Orive 1995; Pineda-Krch & Lehtila 2002). These models are supported by Yu et al. (2020), who identified thousands of single nucleotide polymorphisms among ramets (individual stems) of common eelgrass (*Zostera marina*) that were impacted by natural selection. Furthermore, Cruzan et al. (2022) observed that seep monkeyflower (*Mimulus guttatus*) exhibited extraordinary variation in fitness due to the accumulation of somatic mutations during stem growth, which in some cases led to higher fitness from potentially beneficial somatic mutations being inherited by progeny. This increased fitness may be a result of the novel environments that the plants were grown in (i.e., salt stress), as somatic mutations that were transmitted to offspring would have a high probability of being beneficial (Fisher 1930). These results suggest that somatic mutations can play a non-negligible—and possibly beneficial—role in plant fitness, challenging earlier studies on the topic, which have claimed that beneficial mutations should be exceedingly rare (Charlesworth & Willis 2009).

To shed additional light on the evolutionary consequences of somatic mutations, we performed multiple experiments to determine the fitness effects of inherited somatic mutations in the bush monkeyflower (*Mimulus aurantiacus* Curtis; Phrymaceae). *M. aurantiacus* is a woody, perennial subshrub that is found throughout semi-arid regions of southwestern North America (McMinn 1951). To track the fitness effects of somatic mutations that accumulate within a single generation, we take advantage of the fact that these shrubs have separate stems. Each stem can thus contain distinct germ cell lineages that are derived from the same zygote. As a consequence, each stem can potentially contain different sets of somatic mutations that have accumulated during growth.

By making crosses either within the same flower (autogamy) or between flowers on separate stems of the same plant (inter-stem geitonogamy—hereafter, just geitonogamy), we can produce progeny segregating for somatic mutations that vary among stems. Critically, these crosses are both self-fertilizations, which leads to high homozygosity of meiotic mutants. However, the offspring of each cross type will differ in the complement of somatic mutations that they inherit. For a diploid plant, we can assume that somatic mutations (*a* → *a′*) will be in the heterozygous state when they first appear. For progeny generated via autogamy, a somatic mutation will segregate as 25% homozygous (*a′a′*), 50% heterozygous (*aa′*), and 25% the original (wildtype) homozygote (*aa*). By contrast, progeny from geitonogamous crosses will segregate for somatic mutations that are different in each stem, such that 50% of offspring will be carrying mutations in the heterozygous state and none of the progeny will be homozygous for mutations that arose in a single stem. Thus, the average fitness effects of somatic mutations can be evaluated by comparing the difference in fitness of progeny generated by autogamous and geitonogamous crosses (Bobiwash et al. 2013; Schultz and Scofield 2009).

As noted above, prior studies in the herbaceous perennial *M. guttatus* demonstrated substantial fitness consequences of somatic mutations when grown under salt stress (Cruzan et al 2022). By investigating the fitness effects of somatic mutations in a large and long-lived, woody shrub (*M. aurantiacus*), we are able to compare results between two closely related plant species that differ in important life history characteristics. Moreover, rather than testing progeny in a novel environment, we challenged progeny under drought conditions - a stress that *M. aurantiacus* routinely encounters in its native habitat (Sobel et al 2019). We followed fitness among these two sets of progeny across multiple stages in the life cycle, including fecundity, germination, early seedling growth rates, survival under terminal drought conditions, and total biomass. Under a model where CLS sieves out deleterious somatic mutations while retaining beneficial ones, we expect to find significant differences in fitness between progeny generated from autogamous and geitonogamous pollination (Cruzan et al 2022). This difference in fitness would be attributable to the accumulation of somatic mutations in vegetative tissue that were subsequently transmitted to progeny. In addition, due to the different patterns of segregation of somatic mutations between cross types, we expect progeny from autogamous pollinations to display increased variation in fitness compared to progeny from geitonogamous crosses (Cruzan et al. 2022). Findings from this study contribute to our understanding of the relevance of somatic mutations in plant evolution.

## Materials and Methods

### Experimental setup

To estimate the fitness effects of somatic mutations, we made autogamous and geitonogamous crosses in 26 *M. aurantiacus* genets that had been growing in an open plot in Eugene, Oregon for four years. These genets were initially created through the crossbreeding of red- and yellow-flowered ecotypes of *M. aurantiacus* ssp. *puniceus* (Sobel & Streisfeld 2015; Chase et al. 2017). Using saturating pollen loads from a single flower at the end of each stem, we made four crosses: one autogamous pollination on that same flower and three geitonogamous pollinations to flowers on different stems of the same genet. Hereafter, we refer to the offspring from a set of autogamous and geitonogamous crosses made from a single pollen donor as a “unit.” Because somatic mutations can arise uniquely in any stem, we created multiple units from different stems on the same genet (mean: 1.8 per genet; range 1 - 4). Specifically, between 1 and 22 July 2021, we made 170 crosses, of which 163 developed into fruits. This included 42 individual units that successfully produced a fruit from the autogamous cross and at least two of the geitonogamous crosses. These were used in subsequent analyses.

We note that this approach is an improvement over the method used in Cruzan et al. (2022), where pollen from two stems was reciprocally crossed to create autogamous and geitonogamous pollinations. In that case, distinct somatic mutations in each stem could not be controlled for, which may have impacted estimates of fitness. By using pollen from a single flower to produce multiple geitonogamous crosses on different stems, we were better able to control for different mutations among stems.

#### Fecundity

Fruits were collected when they turned brown and stored at room temperature for two months to allow them to mature fully. Each mature fruit was weighed to the nearest 0.1 mg. Seeds were carefully separated from their capsule, and all seeds from each fruit were weighed. Seeds were then photographed using a Sony Alpha 6000 digital camera and counted using ImageJ software.

### Seed germination and growth rate

From four units (two units each from genets A and C), we performed a germination experiment to determine if the time to germinate differed between pollination treatments. For each of the four units, we filled two 96-cell plug trays with moist potting soil and randomly sowed 192 seeds derived from autogamy and 192 seeds from geitonogamy (64 seeds from each of the three geitonogamous crosses) across the cells (two seeds of the same cross type per cell). Trays were placed in a grow room equipped with fluorescent lights and maintained at 22C with a 16-hour photoperiod. Trays were bottom-watered and overhead misted as needed. Seedling emergence was recorded at the same time each day for 16 days after the first seedling emerged. Each day, seedlings were digitally photographed from above with a ruler in the frame, and we estimated total leaf area using Adobe Photoshop. To estimate early seedling growth rates, we subtracted the total leaf area on the first day a seedling emerged from the total leaf area on the final day of the experiment and divided this by the number of days since the seeding emerged.

### Drought Sensitivity

In the Cruzan et al (2022) study, the fitness of *M. guttatus* offspring was measured in a novel greenhouse environment. However, the ecotypes of *M. aurantiacus* ssp. *puniceus* are drought tolerant shrubs that have adapted to endure seasonal droughts in southern California (Sobel et al. 2019). Because of these seasonal droughts, drought sensitivity likely serves as a principal agent of selection for these ecotypes in the wild. Therefore, to determine if somatic mutations can impact the fitness of offspring under drought conditions, we employed a terminal drought experiment (as in Sobel et al 2019).

Using the seedlings from the germination experiment, we randomly selected 48 plants from autogamous crosses and 48 seedlings from geitonogamous crosses (16 per cross) to transplant into individual cone-shaped pots (21 cm deep) filled with potting soil, which were placed into random positions within a separate 98-cell rack for each unit. Racks were placed in the University of Oregon greenhouse and bottom watered as needed for two weeks to allow seedlings to recover and to establish their roots in the deep cones. After this, no water was added. On each subsequent day, a single researcher categorically scored plant health using a scale between 0 and 4 (as described in Sobel et al. 2019). A score of 0 indicated no sign of drought stress. A score of 1 indicated initial signs of drought stress, including the adaxial side of the leaves curling under. A score of 2 indicated the first sign of true wilting. A score of 3 indicated systemic and severe wilting. A score of 4 indicated death of the plant. The experiment ended once all plants were assigned a score of 4. Plants were measured at the same time each day throughout the experiment, and the identity of the pollination treatment was kept blind to the evaluator until the end of the experiment. At the end of the experiment, the above ground plant material was harvested, dried, and weighed to provide a final estimate of biomass at the time the plant died. To test the effects of somatic mutations across a broader set of stems, we repeated the drought experiment using six additional units (one unit from each of six additional genets; a total of 960 seedlings measured among the 10 units), but we did not collect germination, growth rate, or biomass data from these plants.

To provide an estimate of drought tolerance from these time-series data, we fit a three-parameter logistic curve to the drought scores estimated in each plant on each day of the experiment. Then, we estimated the parameter ‘*b*,’ which occurs at the time (in days) when the drought score reaches 50% of its maximum. This corresponds to the rate at which each plant begins showing obvious signs of drought stress, such that a larger value of ‘*b*’ indicates a more drought tolerant plant. This was repeated separately for each plant within each of the 10 units used in the drought experiments. We also estimated the time (in days) for plants to reach a drought score of 4 (i.e., the survival time). Prior to analysis, we removed 11 plants that died too quickly to obtain accurate parameter estimates.

### Data Analysis

Our primary goal was to determine if there were fitness differences between offspring generated from autogamy and geitonogamy. To begin, we averaged the seed counts and seed and fruit weights from the multiple geitonogamous crosses per unit and used separate paired t-tests to determine if fruit weight, seed weight, and seed count differed significantly between autogamous and geitonogamous pollinations. We then standardized the individual fitness components from each unit to a mean of zero and standard deviation of one. We performed separate MANOVAs for each unit to test if the five fitness components estimated on each seedling differed between pollination treatments. Statistical significance was tested using Pillai’s trace, and effect size was calculated using the partial eta-squared method (Cohen 1988). Individual linear models were then performed with each fitness component as the response variable and cross type as the predictor variable in each of the four units to determine which aspects of fitness differed between pollination type. Finally, we estimated the coefficient of variation between autogamous and geitonogamous treatments for each fitness component across the four units to determine if the variance in fitness was higher in progeny from autogamous crosses, as predicted under a model of cell lineage selection. All analyses were performed in R.

## Results

### Fecundity

We identified significant effects of cross type on fecundity. Specifically, among the 42 units from 26 genets generated in this experiment, autogamous pollination consistently resulted in more seeds than geitonogamous pollination (paired t-test, p = 0.014; Fig 1). Indeed, in 29 of the 42 units (69%), the total seed count per fruit was higher from autogamous crosses. This pattern was similar for fruit weight and total seed weight as well (both p = 0.004), which were each strongly correlated with seed count (seed count vs seed weight: *r* = 0.87; seed count vs fruit weight: *r* = 0.58).

**Figure 1.**
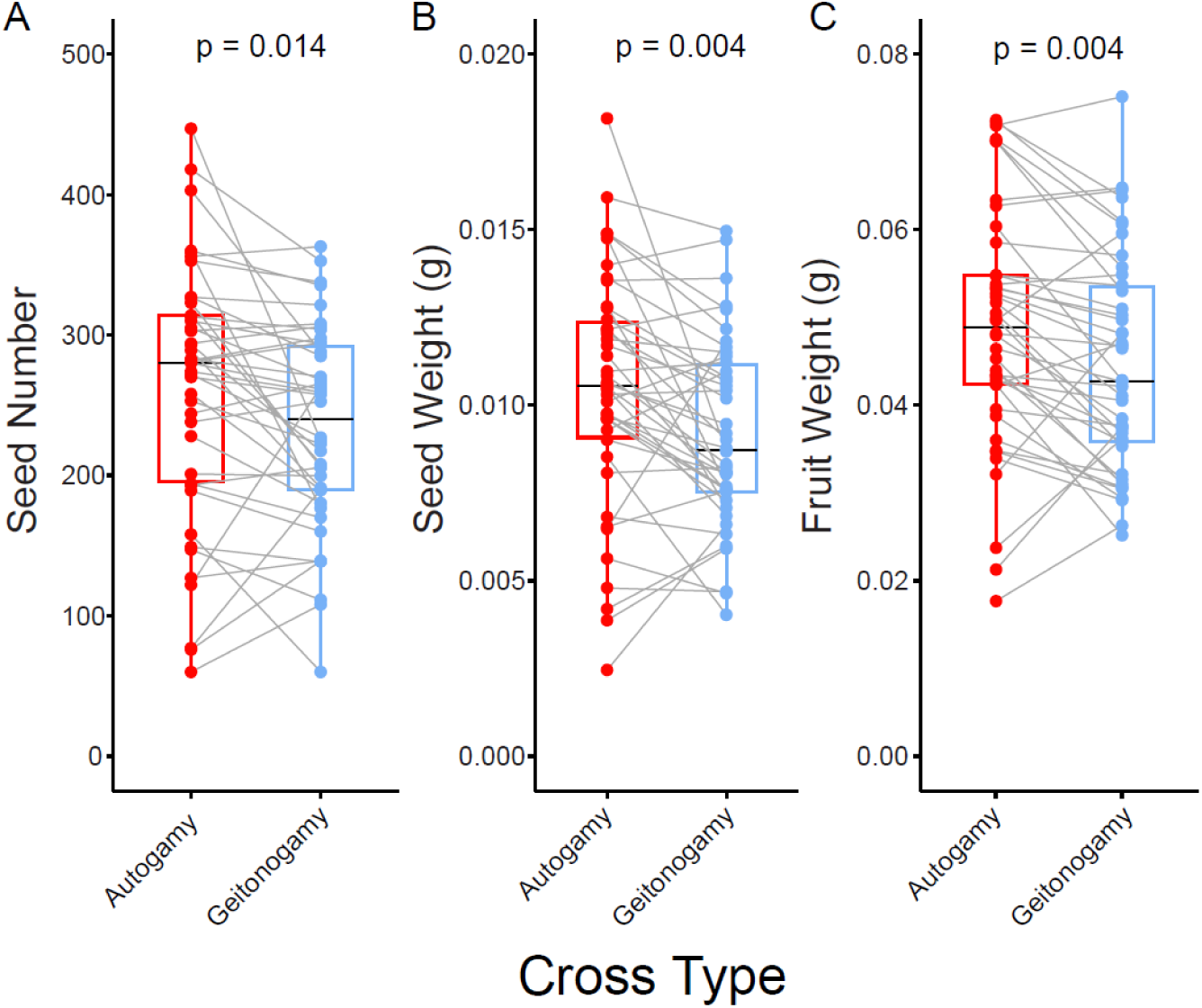
Fecundity is higher following autogamous pollination compared to geitonogamous pollination. In each panel, gray lines connect the fecundity estimates from autogamous and geitonogamous pollinations from each of the 42 units in the experiment. Box plots show the median (in black), the bottom and top of the boxes correspond to the first and third quartile, respectively, and whiskers represent 1.5 times the interquartile range. P-values above each plot were estimated using paired t-tests. A) Seed number per fruit, B) total seed weight, C) total fruit and seed weight. Values for geitonogamous crosses were averaged from the two or three crosses made within each unit.

### Patterns of selection in offspring

We measured variation in five aspects of fitness among the offspring of autogamous and geitonogamous pollinations. These components of selection acted at different stages of the plant life cycle, beginning with germination, and continuing through early seedling growth rates, drought tolerance, survival, and total biomass. Using MANOVA, we found an overall significant difference in fitness for progeny derived from autogamous and geitonogamous crosses in two of the four units (A1 and C2; Table S1). In both cases, the partial eta-squared value > 0.14, indicating a moderate to large effect of cross type on the multivariate fitness estimates (Cohen 1988). By contrast, the other two units (A2 and C1), which were derived from different stems of these same two genets, showed no difference between pollination types (P > 0.155). These results demonstrate variation in fitness among stems of the same genet, likely due to different complements of somatic mutations having accumulated in each stem.

In the offspring of unit A1, there were significant differences in drought tolerance, survival, and biomass between pollination treatments (Table S2), with the mean fitness being higher in autogamous crosses for drought tolerance and survival and lower for biomass (Fig 2, Fig S1). Differences in drought tolerance were negatively correlated with both early seedling growth rates and biomass (Fig S2, S3), consistent with previous findings in this species that revealed smaller plants were better able to withstand drought conditions (Sobel et al 2019). Interestingly, even though we found an overall significant effect of cross type on multivariate fitness in unit C2, none of the five selection components were individually significant between pollination types (Table S2). This implies that despite an effect of cross type when all components are tested together, that effect is not driven strongly by any one measure of fitness. By contrast, in unit C1, drought tolerance was significantly higher in offspring derived from autogamous crosses (Table S2), even though the overall MANOVA was not significant (Table S1).

**Figure 2.**
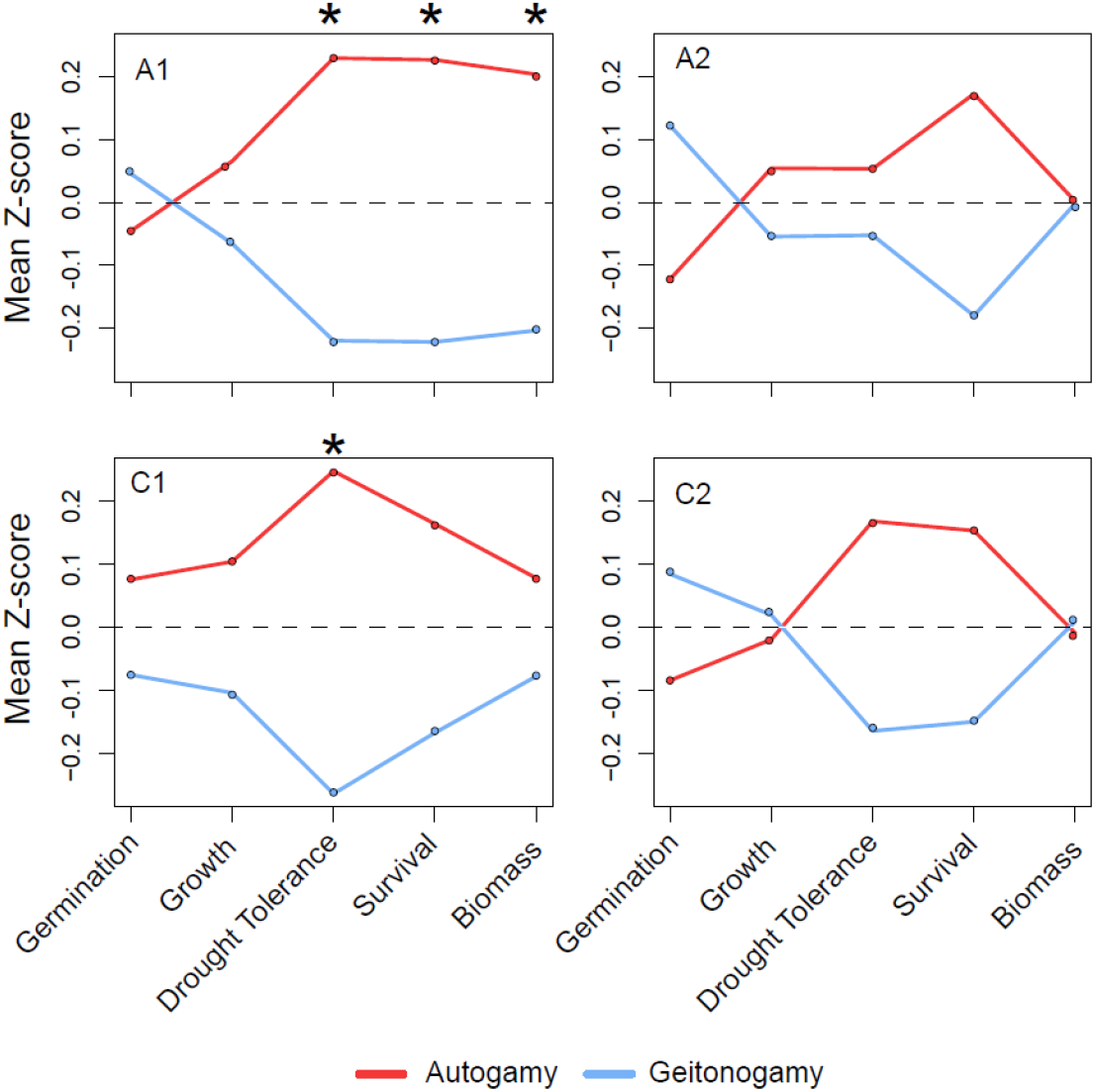
Mean fitness varies between cross types across the four experimental units. The five fitness components are listed at the bottom, in order of their occurrence across the life-cycle. The data are presented as mean Z-scores for each fitness component broken down by cross type, such that the overall mean is 0 and the standard deviation is 1. The gray dashed line is at zero, corresponding to the expected mean values across both pollination treatments. Values above and below zero correspond to the number of standard deviations above and below the mean, respectively. Individual boxplots of each fitness component are presented in Fig. S1. To aid in visual presentation, individual Z-scores for growth rate and biomass were inverted by multiplying values by -1, and the mean was estimated. Asterisks indicate statistically significant differences between pollination treatments (p < 0.05).

**Figure 3.**
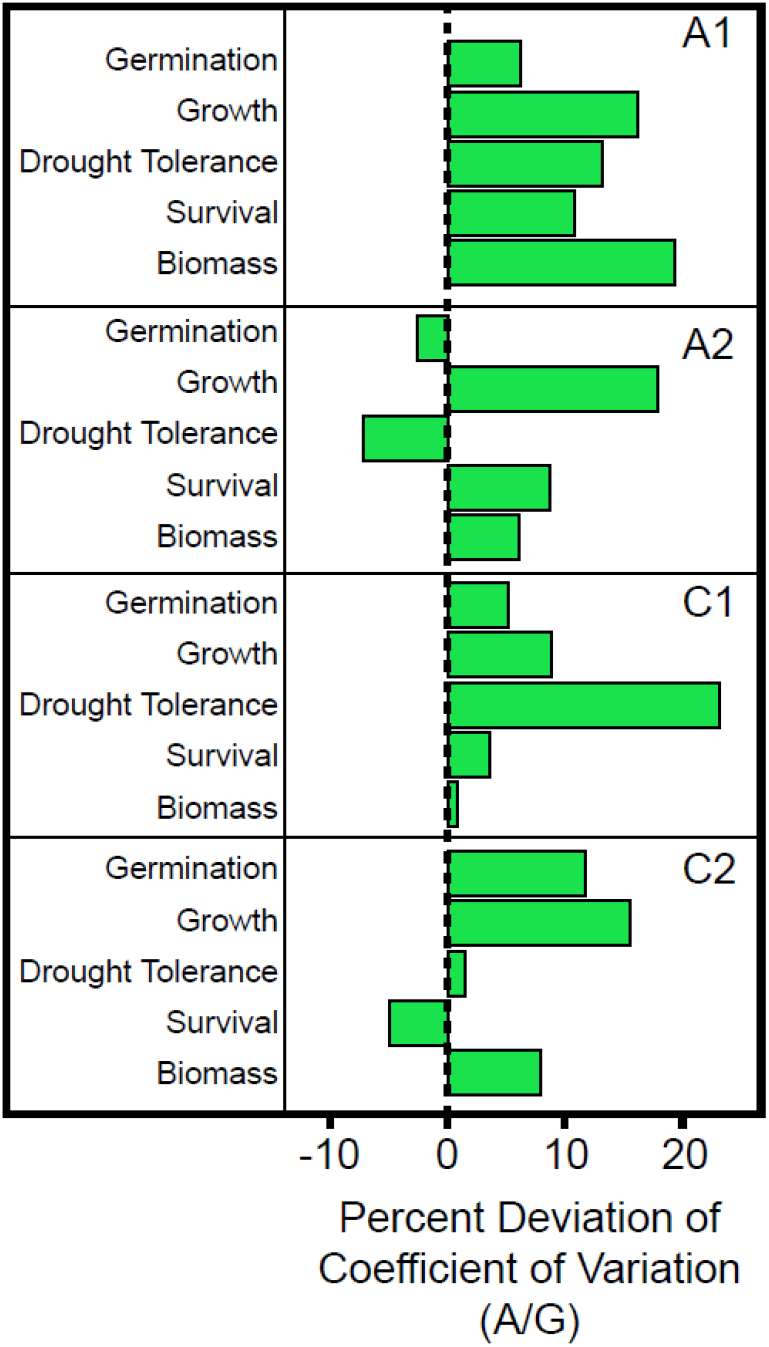
The variance in fitness in offspring is higher following autogamy compared to geitonogamy. Shown is the percent deviation in the coefficient of variation between autogamous and geitonogamous pollination treatments for the different finess estimates across all four units. A value of zero would indicate that the coefficient of variation for a particular fitness estimate is the same between autogamy and geitonogamy, but positive values reveal higher variation in fitness among offspring derived from autogamy compared to geitonogamy.

Across all estimated fitness components, we did not find an effect of cross type on the timing of germination or early seedling growth rates. However, we did observe differences in drought tolerance in two of the four units. Therefore, we tested whether this pattern was consistent among a larger set of units from six additional genets. In one of the six units, there was a significant difference in drought tolerance between cross types, with plants derived from geitonogamous pollination having a slightly higher mean value of drought tolerance (Table S3). There were no differences between cross types in the other five units.

In addition to differences in fitness, we also predicted that selection occurring in progeny following the transmission of somatic mutations that accumulated in stems would result in a higher variance in fitness in the offspring from autogamous pollinations compared to geitonogamous pollinations. If somatic mutations affect offspring fitness, variation in fitness should be greater for progeny groups from autogamy than from geitonogamy, as long as mutations are not completely dominant. This is because somatic mutations will segregate as homozygotes and heterozygotes in autogamous progeny but will remain heterozygous in the progeny of geitonogamous crosses. To investigate this, we compared the coefficient of variation for each of the five fitness components between autogamous and geitonogamous treatments. We find that the coefficient of variation is higher in offspring from autogamy in 17 of the 20 cases, with a deviation that averages 10.9% higher following autogamy than geitonogamy. In particular, in unit C1, we see that the variation in drought tolerance is 23.2% higher in the autogamy treatment compared to geitonogamy. Similarly, we see higher variance among autogamous offspring for drought tolerance, survival, and biomass in unit A1 (range 10.8 - 13.9 %), the three components that were significantly different between pollination treatments. The excess variance in the autogamy treatment compared to geitonogamy is highly significant across all units combined (binomial probability, p = 0.001), implying that there is overall higher variance in fitness following autogamy, consistent with the effects of somatic mutations accumulating within stems and affecting fitness as they segregate in offspring.

## Discussion

In this study, we demonstrated that the accumulation of somatic mutations in vegetative tissue can impact the fitness of plants in the following generation. In addition, rather than somatic mutations being uniformly deleterious, we show that they can occasionally have a net beneficial effect, resulting in an increase of average fitness. This finding is consistent with expectations from models of cell lineage selection (Fagerstrom et al. 1998; Otto and Hastings 1998; Monro and Poore 2009), which argue that cell lineages with faster growth can displace slower ones (Poethig 1987; Klekowski 2003). If these differences in division rates are determined by somatic mutations, we would expect CLS to contribute to the purging of mutational load (Pineda-Krch and Fagerstrom 1999; Monro and Poore 2009). Similarly, we expect mutations enhancing growth to be retained. Therefore, cell lineage selection during vegetative growth has the potential to modify the distribution of fitness effects of accumulated mutations by filtering expressed deleterious mutations and allowing the transmission of beneficial variants. Specifically, we found evidence that supports the accumulation and transmission of somatic mutations, which can lead to higher fecundity, and in some cases, increased tolerance under drought conditions. These results provide evidence for the potential importance of somatic mutations for plant evolution.

In spite of slow division rates and possibly enhanced DNA repair capacity (Yadav et al. 2009; Heyman et al. 2013), plant meristem cells are expected to accumulate substantial levels of mutational load during stem elongation. The effects of this deleterious variation often are apparent as reduced fecundity (or increased embryo abortion) following autogamous compared to geitonogamous pollinations, which has been referred to as autogamy depression (Schultz and Scofield 2009). Autogamy depression for seed and fruit abortion has been observed in several species (reviewed in Bobiwash et al. 2013), including *M. guttatus* (Cruzan et al. 2022), and it is expected to be stronger in longer lived plants, as longer lifespan should correspond to more mitotic cell divisions and thus a greater opportunity for somatic mutation accumulation (Schultz and Scofield 2009, Ally et al. 2010, Barrett 2015). Although *M. aurantiacus* is a long-lived perennial shrub, we did not find evidence for autogamy depression. By contrast, we found an overall average increase in fecundity following autogamy. Given that both cross types are self-fertilizations, these differences cannot be attributable to variation in the strength of inbreeding depression between treatments. Rather, the absence of autogamy depression in this system could be due to the transmission of beneficial somatic variants whose fitness effects outweigh those of deleterious mutations, resulting in a net increase in fecundity. Because deleterious mutations can be filtered out due to CLS prior to fertilization, the presumed larger number of mitotic divisions in these plants may actually result in a shift in the distribution of fitness effects that is skewed toward the transmission of more beneficial mutations rather than deleterious ones. Although these findings conflict with trends seen in other species that show autogamy depression is common (Klekowski 1998, Bobiwash et al 2013), they are consistent with the pattern of an unexpectedly high transmission of beneficial mutations in mutation accumulation studies in *Arabidopsis thaliana* (Shaw et al. 2002; Rutter et al. 2010; Rutter et al. 2012; Rutter et al. 2018). Moreover, it is important to note that the study of the fitness consequences of somatic mutations is in its infancy. Therefore, further investigation on the consistency of these patterns among closely related plants with different life history strategies is needed.

In addition to fecundity, we also measured variation in five aspects of fitness among the offspring of autogamous and geitonogamous pollinations. These components of selection acted at different stages of the plant life cycle, beginning with germination, and continued through early seedling growth rates, drought tolerance, survival, and total biomass. While we did not find significant differences in fitness between pollination treatments for germination or early seedling growth rates in any of the units, we did find evidence for increased tolerance to drought, higher survival, and lower total biomass in offspring derived from autogamy. We also found higher variance in fitness among offspring derived from autogamy, which is in line with our results demonstrating an increase in fitness in seedlings following autogamous pollination (Cruzan et al 2022). In one case (unit E3), we also found significantly higher fitness in seedlings following geitonogamy, which suggests that the transmission of deleterious somatic mutations may have occurred in the autogamous lines. Regardless, these results are consistent with the hypothesis that somatic variants that accumulated during vegetative growth can be transmitted to offspring where they can occasionally impact fitness. Observed increases in fitness after autogamy suggest a potential role for somatic variation in local adaptation.

Despite detecting significant differences for fitness components in some of the units, the individual effect sizes are rather small. This result is not unexpected for at least two reasons.

First, offspring were grown in an environment that closely matches their native habitat. The populations of *M. aurantiacus* used in this experiment occur in chapparal communities of southern California, which is dominated by hot, dry summers and cool, moist winters (Beeks 1962). Seedling recruitment tends to be very low due to the rapid drying of the soil after seedlings emerge. Thus, the terminal drought experiment we conducted closely mimics the conditions of natural seedlings (Sobel et al 2019). As a result, we would expect plants to already be near their adaptive peaks for drought tolerance, suggesting that most new mutations would not greatly improve fitness (Orr 2005). By contrast, previous work in the herbaceous *M. guttatus* revealed that somatic mutations accumulating during vegetative growth had large, beneficial effects on offspring fitness in five of the 14 stems tested (Cruzan et al 2022). In this case, the progeny were grown in a novel environment (hydroponic salt-stress), implying that there was a broader spectrum of mutations that could have phenotypic effects capable of moving the population closer to its optimum. Second, our analyses focused on testing for average differences in fitness between pollination treatments. Following autogamous pollination, only 25% of offspring on average are expected to be homozygous for a somatic mutation that arose in that stem. Therefore, provided that new mutations are not completely dominant, most somatic variants will fail to be expressed in offspring, resulting in few plants that show differences in fitness between cross types. As a consequence, our findings are consistent with a prediction of small differences in average fitness between pollination treatments.

Although the segregation of somatic mutations in offspring can obscure overall statistical patterns between pollination treatments, we can still see the net fitness effects of these variants in individual progeny. Specifically, we observed individual plants derived from autogamous pollination that have exceptional values of fitness, especially for drought tolerance, survival, and biomass. For example, in units A1 and C1 (the units that show significant differences in drought tolerance between pollination treatments), we see that the plants with the highest drought tolerance are derived from autogamy. These plants have drought tolerance values that are more than three standard deviations above the mean (Fig S4). This trend continues with the later-acting fitness components, such that these same plants also have consistently extreme values of survival and biomass. We also found a strong, negative relationship between drought tolerance and early seedling growth rate, such that smaller plants tended to survive longer under drought conditions. These results are consistent with those of Sobel et al (2019), who also found that smaller *M. aurantiacus* plants tended to better withstand desiccation. They suggested that the reduced leaf area of smaller plants likely resulted in lower transpiration, leading to greater drought tolerance and thus longer survival under terminal drought conditions. Thus, the segregation of somatic variants can result in progeny with extreme values of fitness, providing an additional source of genetic variation for adaptation. Future experiments that take advantage of the power of deep sequencing can be used to identify individual somatic variants that accumulated in parents, which would allow us to track the fitness consequences of these variants after they are transmitted to offspring.

In conclusion, we find evidence for the transmission of both beneficial and deleterious somatic variation in offspring, revealing that somatic variation can occasionally underlie adaptation. By comparing these results with those from the closely related *M. guttatus* with different life history characteristics (Cruzan et al 2022), we found similar, though more subtle, fitness consequences following autogamy. Furthermore, as noted earlier, the current study improved on the crossing design used by Cruzan et al (2022). In this case, we used pollen from the same flower for both an autogamous pollination, as well as multiple geitonogamous crosses on different stems of the same genet, which allowed us to control for somatic variation among stems. Thus, the fact that our results are consistent with those from Cruzan et al (2022), despite differences in crossing design, environmental conditions, and life history, reveals the potential relevance of somatic mutation for plant evolution.

Considering that the accumulation and transmission of somatic mutations may be a general feature of plant evolution can provide some insight into the success and diversification of flowering plants. The evolution of apical meristems and indeterminate growth in early land plants may have influenced the potential for cell lineage selection to affect the distribution of mutations acquired during vegetative growth. While the primary selective advantage for producing reproductive structures at the ends of growing stems may have been for improved dispersal, this architecture also maximized the potential for selection among cell lineages to affect the distribution of mutations passed on to offspring. Even though plants are sedentary over much of their life cycle and may be subjected to substantial environmental variation within a single lifespan, we show here that the accumulation of somatic variation during vegetative growth has the potential to contribute significantly to plant adaptation in subsequent generations. Future work in population genetics should not ignore somatic mutations as an important source of genetic variation that can impact plant evolution.

## Acknowledgments

We would like to thank members of the Cruzan lab for their feedback on earlier drafts of this manuscript. We also thank S. Medbury for his assistance with plant care in the University of Oregon greenhouses. This work was supported by NSF-DEB 2051242 to MAS and MBC.

## Supplemental Material

**Table S1.** Results from the MANOVAs for each unit, testing the combined effects of the five fitness components by pollination treatment.

| Unit | Pillai | Eta-squared | P |
| --- | --- | --- | --- |
| A1 | 0.16865 | 0.17 | 0.0055 |
| A2 | 0.08913 | 0.09 | 0.1522 |
| C1 | 0.07724 | 0.08 | 0.2124 |
| C2 | 0.1358 | 0.14 | 0.0215 |

**Table S2.** Results from linear models testing each fitness estimate against pollination treatment for each of the four units. The F-value of the test, degrees of freedom (Df) and the P-value are reported. P-values less than 0.05 are in bold.

| Unit | Fitness estimate | F | Df | P |
| --- | --- | --- | --- | --- |
| A1 | Germination | 0.197 | 1, 94 | 0.659 |
| A1 | Growth rate | 0.409 | 1, 94 | 0.524 |
| **A1** | **Drought tolerance** | **4.937** | **1, 92** | **0.029** |
| **A1** | **Survival** | **4.975** | **1, 93** | **0.028** |
| **A1** | **Biomass** | **4.103** | **1, 94** | **0.046** |
| A2 | Germination | 1.440 | 1, 94 | 0.233 |
| A2 | Growth rate | 0.281 | 1, 94 | 0.597 |
| A2 | Drought tolerance | 0.251 | 1, 89 | 0.617 |
| A2 | Survival | 2.980 | 1, 92 | 0.088 |
| A2 | Biomass | 0.002 | 1, 94 | 0.988 |
| C1 | Germination | 0.548 | 1, 94 | 0.461 |
| C1 | Growth rate | 1.043 | 1, 94 | 0.310 |
| **C1** | **Drought tolerance** | **6.464** | **1, 91** | **0.012** |
| C1 | Survival | 2.665 | 1, 93 | 0.106 |
| C1 | Biomass | 0.582 | 1, 94 | 0.447 |
| C2 | Germination | 0.684 | 1, 94 | 0.410 |
| C2 | Growth rate | 0.036 | 1, 94 | 0.851 |
| C2 | Drought tolerance | 2.684 | 1, 93 | 0.105 |
| C2 | Survival | 2.216 | 1, 93 | 0.140 |
| C2 | Biomass | 0.008 | 1, 94 | 0.927 |

**Table S3.** Results from linear models testing the effects of pollination treatment on drought tolerance in six additional units. The F-value of the test, degrees of freedom (Df) and the P-value are reported. P-values less than 0.05 are in bold.

| Unit | F | Df | P |
| --- | --- | --- | --- |
| **E3** | **6.312** | **1, 94** | **0.0137** |
| F1 | 1.01 | 1, 94 | 0.3175 |
| H1 | 0.295 | 1, 94 | 0.5883 |
| K1 | 0.112 | 1, 94 | 0.7382 |
| O1 | 0.081 | 1, 94 | 0.7765 |
| R2 | 0.396 | 1, 94 | 0.5308 |

**Figure S1.**
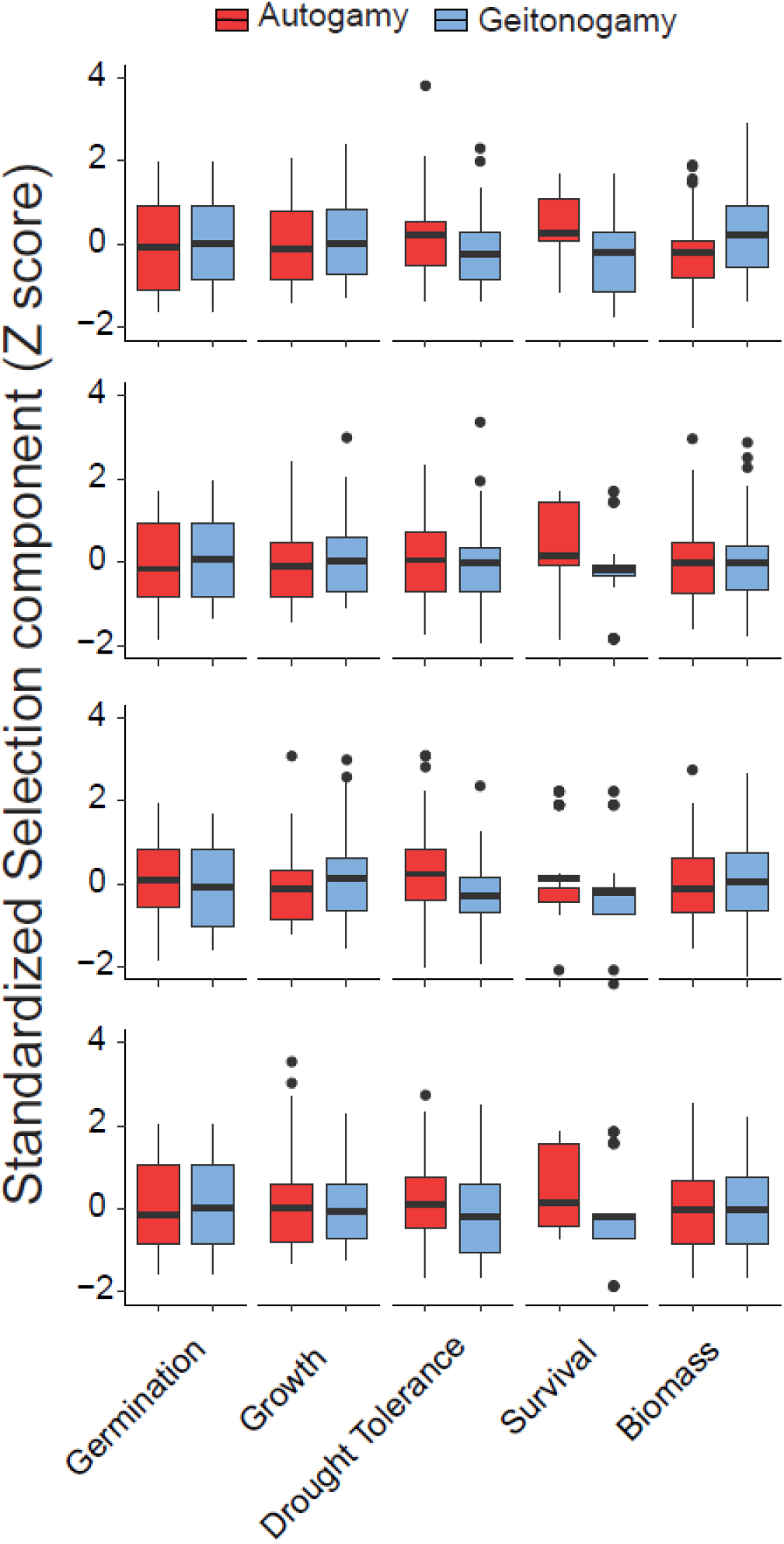
Boxplots of the fitness estimates for each fitness component across the four units following autogamy (red) and geitonogamy (blue). Fitness values are standardized to z-scores, with a mean of zero and standard deviation of 1. The black horizontal line corresponds to the median, box heights indicate the lower and upper quartile, and whiskers correspond to 1.5 times the interquartile range. From top to bottom, plots are for units A1, A2, C1, C2.

**Figure S2.**
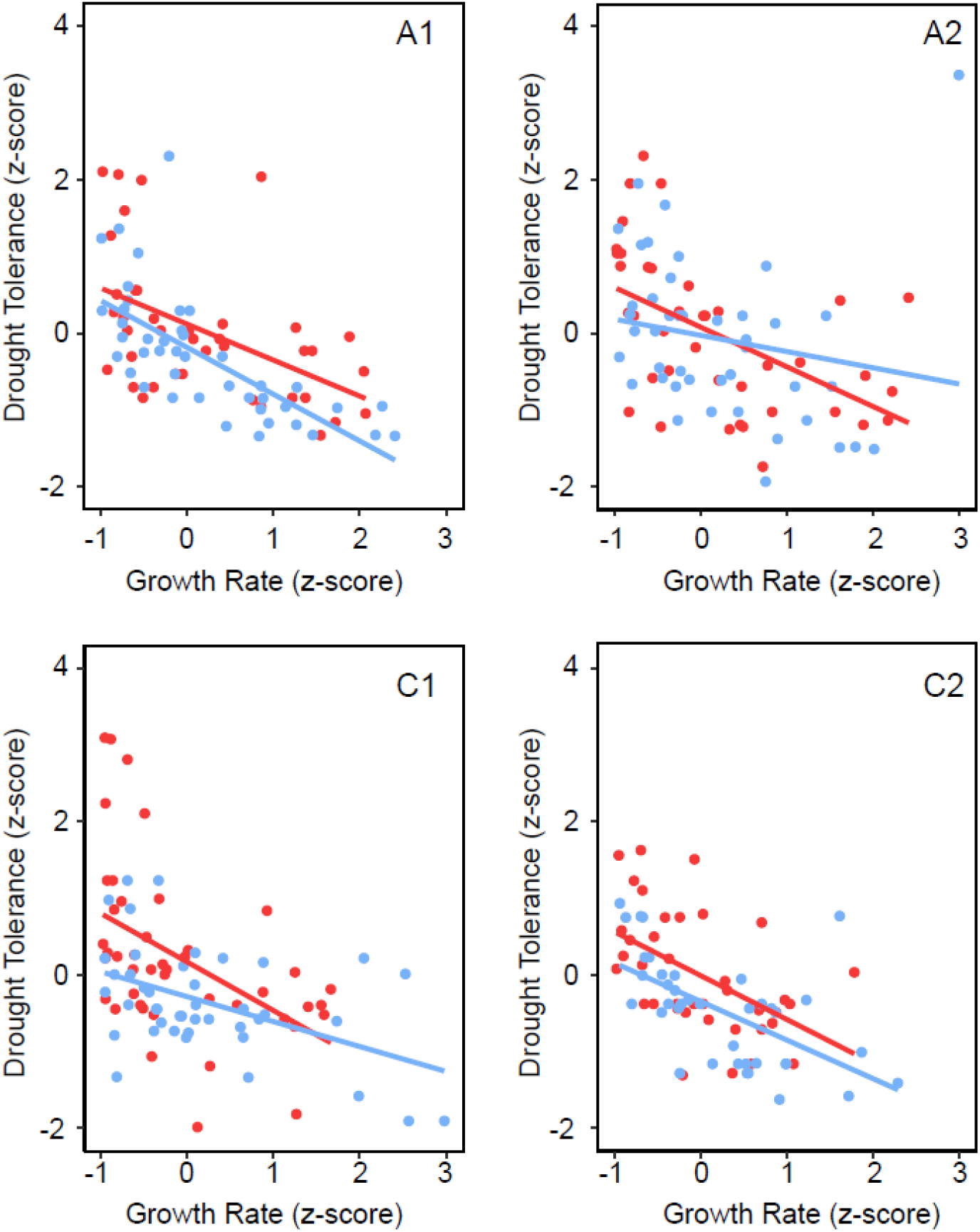
The relationship between drought tolerance and early seedling growth rate across the four units. Red points correspond to seedlings derived from autogamy and blue points correspond to seedlings derived from geitonogamy. Trendlines are derived from linear models testing the effect of drought tolerance against growth rate, separately for each pollination treatment. Fitness values are standardized to z-scores, with a mean of zero and standard deviation of 1.

**Figure S3.**
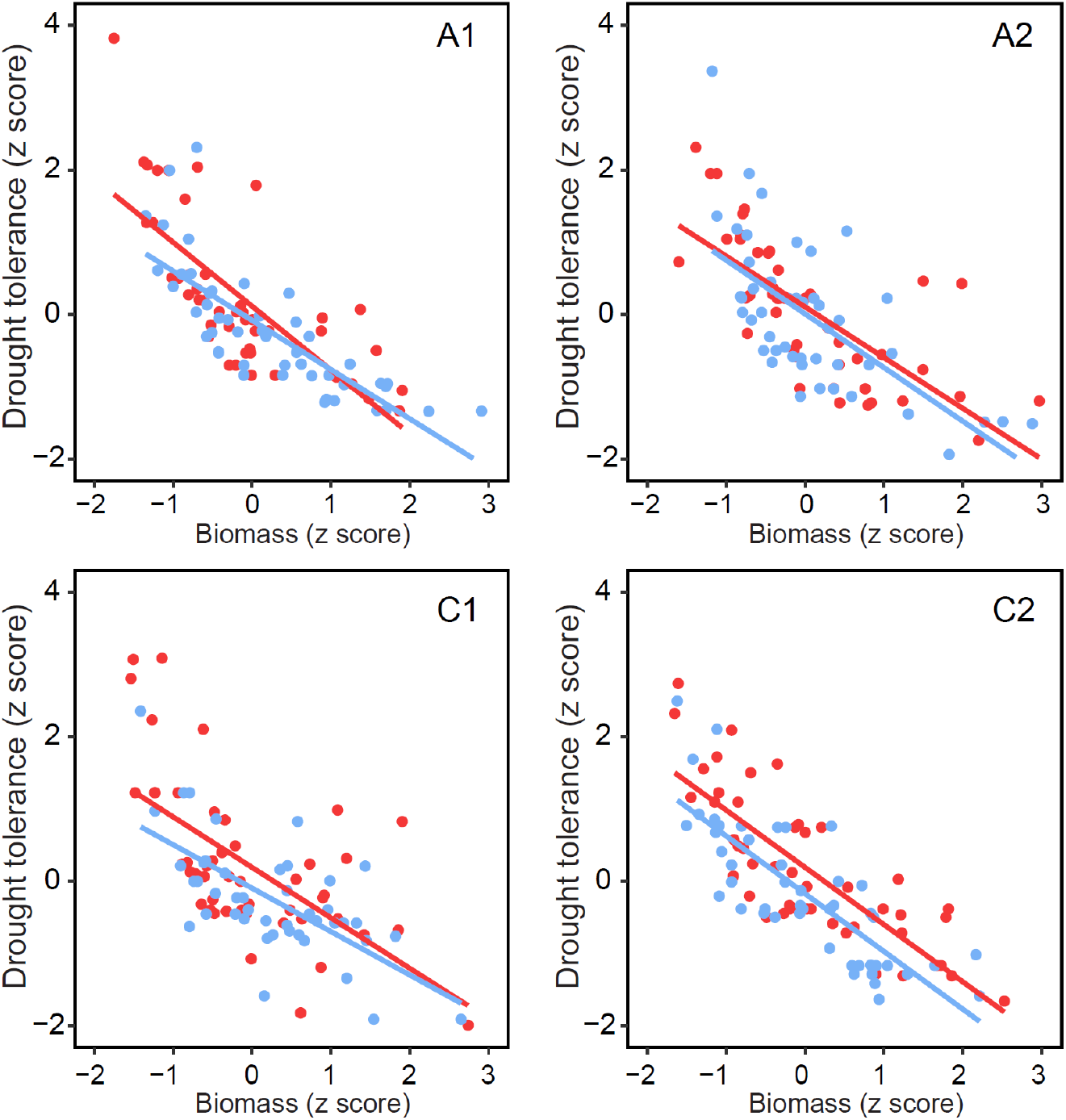
The relationship between drought tolerance and biomass across the four units. Red points correspond to seedlings derived from autogamy and blue points correspond to seedlings derived from geitonogamy. Trendlines are derived from linear models testing the effect of drought tolerance against biomass, separately for each pollination treatment. Fitness values are standardized to z-scores, with a mean of zero and standard deviation of 1.

**Figure S4.**
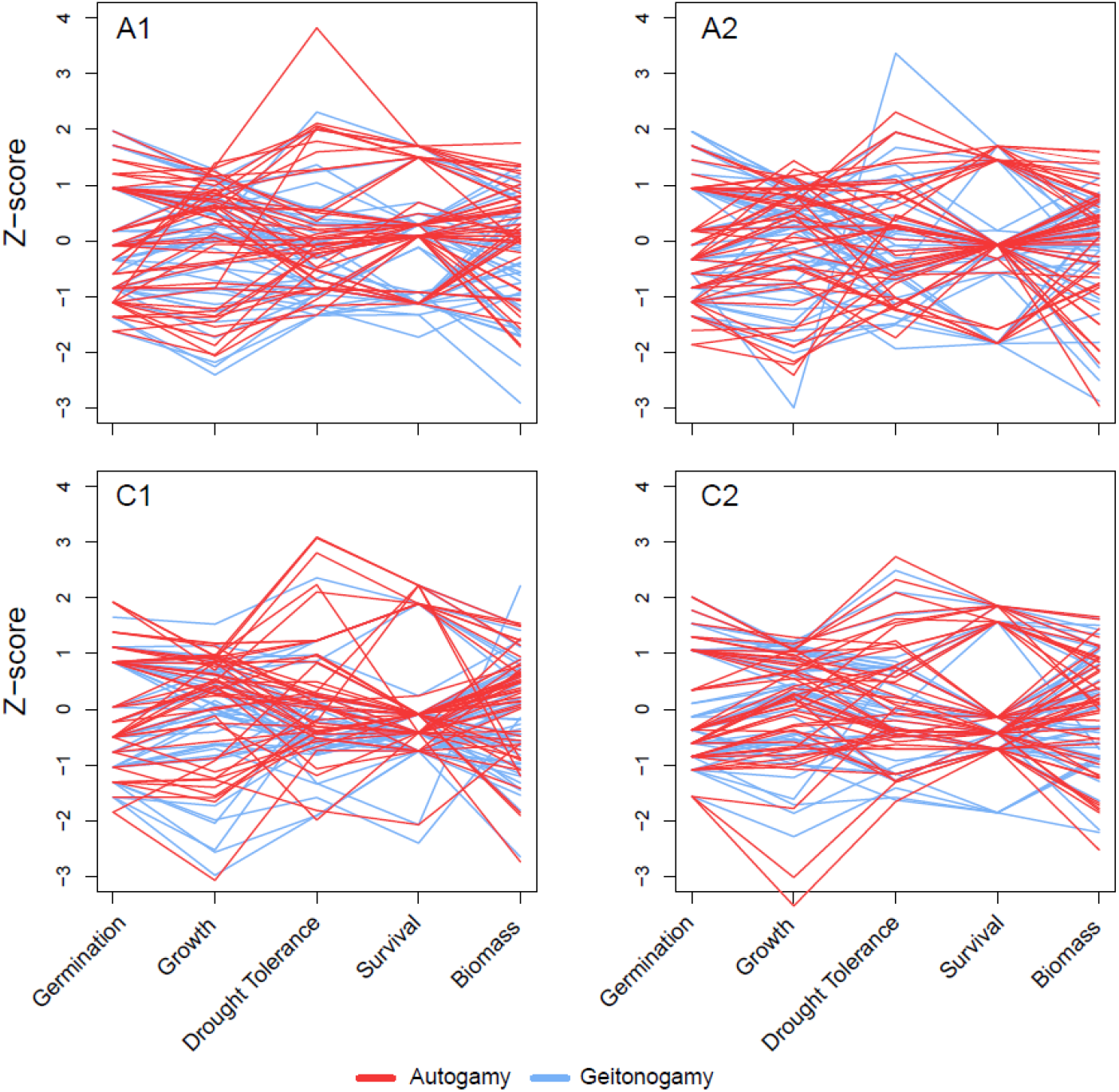
Line plots connecting fitness estimates for individual seedlings derived from either autogamy (red) or geitonogamy (blue) across the four units. Fitness values are standardized to z-scores, with a mean of zero and standard deviation of 1.

## Notes

### Competing Interest Statement

The authors have declared no competing interest.

## References

Ally, D., K. Ritland, and S. P. Otto. 2010. Aging in a Long-Lived Clonal Tree. PLoS Biol. 8.

Antolin, M. F. & Strobeck, C. (1985). The population genetics of somatic mutation in plants. Am. Nat. 126:52–62.

Barrett, S. C. H. 2015. Influences of clonality on plant sexual reproduction. Proceedings of the National Academy of Sciences 112:8859–8866

Beeks, R. M. 1962. variation and hybridization in southern California populations of Diplacus (Scrophulariaceae). El Aliso 5:83–122.

Bobiwash, K., S. T. Schultz, and D. J. Schoen. 2013. Somatic deleterious mutation rate in a woody plant: estimation from phenotypic data. Heredity 111:338–344.

Burian, A. & Barbier de Reuille, K. (2016). Patterns of stem cell divisions contribute to plant longevity. Current Biology. 26(11):1385–1394.

Charlesworth, D. & Willis, J. H. (2009). The genetics of inbreeding depression. Nature Reviews Genetics. 10: 783–796.

Chase, M. A., S. Stankowski, and M. A. Streisfeld. 2017. Genomewide variation provides insight into evolutionary relationships in a monkeyflower species complex (Mimulus sect. Diplacus). American Journal of Botany 104:1510–1521.

Cohen, J (1988) Statistical power analysis for the behavioral sciences (2nd ed.). Hillsdale, NJ: Erlbaum.

Cruzan, M. B. 2018. Evolutionary Biology – A Plant Perspective. Oxford University Press, New York.

Cruzan, M. B., M. A. Streisfeld, and J. A. Schwoch. 2022. Fitness effects of somatic mutations accumulating during vegetative growth. Evol. Ecol. 36:767–785.

Cutter, E. G. (1965). Recent experimental studies of the shoot apex and shoot morphogenesis. Botanical Review. 31(1): 7–21.

D’Amato, F. (1996). Role of somatic mutations in the evolution of higher plants. Caryologia. 50(1).

Fagerstrom, T., D. A. Briscoe, and P. Sunnucks. 1998. Evolution of mitotic cell-lineages in multicellular organisms. Trends Ecol. Evol. 13:117–120.

Fisher, Ronald (1930). The Genetical Theory of Natural Selection. Oxford, UK: Oxford University Press.

Gaut, B., Yang, L., Takuno, S., Eguiarte, L. E. (2011). The patterns and causes of variation in plant nucleotide substitution rates. Annual review of ecology, evolution, and systematics. 42: 245–266.

Heyman, J., T. Cools, F. Vandenbussche, K. S. Heyndrickx, J. Van Leene, I. Vercauteren, S. Vanderauwera, K. Vandepoele, G. De Jaeger, D. Van Der Straeten, and L. De Veylder. 2013. ERF115 Controls Root Quiescent Center Cell Division and Stem Cell Replenishment. Science 342:860–863.

Klekowski, E. J. 1988. Mutation, Developmental Selection, and Plant Evolution. Columbia University Press, New York.

Klekowski, E. J. 2003. Plant clonality, mutation, diplontic selection and mutational meltdown. Biol. J. Linn. Soc. 79:61–67.

Kwiatkowska, D. (2008). Flowering and apical meristem growth dynamics. Journal of Experimental Botany. 59(2): 187–201.

Lanfear, R. (2018). Do plants have a segregated germline?. PLOS Biology. 16(5).

McMinn, H. E. 1951. Studies in the genus Diplacus, Scrophulariaceae. Madrono 11:33–128.

Monro, K., and A. G. B. Poore. 2009. The Potential for Evolutionary Responses to Cell-Lineage Selection on Growth Form and Its Plasticity in a Red Seaweed. Am. Nat. 173:151–163.

Orr, H. A. 2005 The genetic theory of adaptation: a brief history. Nature Reviews Genetics. 6:119–127.

Otto, S. P., and I. M. Hastings. 1998. Mutation and selection within the individual. Genetica 102–3:507–524.

Otto, S. P., Orive, M. E. (1995). Evolutionary consequences of mutation and selection within an individual. Genetics. 141(3): 1173–1187.

Pineda–Krch, M., and T. Fagerstrom. 1999. On the potential for evolutionary change in meristematic cell lineages through intraorganismal selection. J. Evol. Biol. 12:681–688.

Pineda-Krch, M. P. & Lehtila, K. (2002). Cell lineage dynamics in stratified shoot apical meristems. Journal of Theoretical Botany. 219(4): 495–505.

Poethig, R. S. 1987. Clonal analysis of cell lineage patterns in plant development. Am. J. Bot. 74:581–594.

Rutter, M. T., A. Roles, J. K. Conner, R. G. Shaw, F. H. Shaw, K. Schneeberger, S. Ossowski, D. Weigel, and C. B. Fenster. 2012. Fitness of Arabidopsis thaliana mutation accumulation lines whose spontaneous mutations are known. Evolution 66:2335–2339.

Rutter, M. T., A. J. Roles, and C. B. Fenster. 2018. Quantifying natural seasonal variation in mutation parameters with mutation accumulation lines. Ecology and Evolution 8:5575–5585.

Rutter, M. T., F. H. Shaw, and C. B. Fenster. 2010. Spontaneous mutation parameters for Arabidopsis thaliana measured in the wild. Evolution 64:1825–1835.

Schultz, S. T. & Scofield, D. G. (2009). Mutation accumulation in real branches: fitness assays for genomic deleterious mutation rate and effect in large-statured plants. The American Naturalist. 174:163–175.

Shaw, F. H., C. J. Geyer, and R. G. Shaw. 2002. A comprehensive model of mutations affecting fitness and inferences for Arabidopsis thaliana. Evolution 56:453–463.

Sobel, J. M., Stankowski, S., Streisfeld, M. A. (2019). Variation in ecophysiological traits might contribute to ecogeographic isolation and divergence between parapatric ecotypes of Mimulus aurantiacus. Journal of Evolutionary Biology. 32: 604–618

Sobel, J. M., Streisfeld, M. A. (2015). Strong premating reproductive isolation drives incipient speciation in Mimulus aurantiacus. International Journal of Organic Evolution. 69(2): 447–461.

Yadav, R. K., T. Girke, S. Pasala, M. T. Xie, and V. Reddy. 2009. Gene expression map of the Arabidopsis shoot apical meristem stem cell niche. Proceedings of the National Academy of Sciences of the United States of America 106:4941–4946.

